# Mechanical characterization of regenerating *Hydra* tissue spheres

**DOI:** 10.1101/2023.10.16.562504

**Authors:** Thomas Perros, Anaïs Biquet-Bisquert, Zacchari Ben Meriem, Morgan Delarue, Pierre Joseph, Philippe Marcq, Olivier Cochet-Escartin

**Affiliations:** Universite Claude Bernard Lyon 1, CNRS, Institut Lumière Matière, Villeurbanne, France; Centre de Biologie Structurale, Université de Montpellier-CNRS-INSERM, Montpellier, France; Laboratory for Analysis and Architecture of Systems, Université de Toulouse-CNRS, UPS, INP, Toulouse, France; Physique et Mécanique des Milieux Hétérogènes, PMMH, CNRS, ESPCI Paris, Université PSL, Sorbonne Université, Université Paris Cité, F-75005, Paris, France

## Abstract

*Hydra vulgaris,* long known for its remarkable regenerative capabilities, is also a longstanding source of inspiration for models of spontaneous patterning. Recently, it became clear that early patterning during *Hydra* regeneration is an integrated mechano-chemical process where morphogen dynamics is influenced by tissue mechanics. One roadblock to understand *Hydra* self-organization is our lack of knowledge about the mechanical properties of these organisms. In this paper, we combined microfluidic developments to perform parallelized microaspiration rheological experiments and numerical simulations to characterize these mechanical properties. We found three different behaviors depending on the applied stresses: an elastic response, a visco-elastic one and tissue rupture. Using models of deformable shells, we quantify their Young’s modulus, shear viscosity as well as the critical stresses required to switch between behaviors. Based on these experimental results, we propose a description of the tissue mechanics during normal regeneration. Our results provide a first step towards the development of original mechano-chemical models of patterning grounded in quantitative, experimental data.

**Statement of significance:** *Hydra vulgaris* is a remarkable organism thanks to its regenerative abilities. One can cut this animal into several pieces which will reform a full *Hydra* in a few days. In this process, the pieces have to define a new organizing axis. Recently, researchers have shown that this axis definition is under mechanical control. One roadblock to understand the relationship between tissue mechanics and *Hydra* biology is our lack of knowledge about the mechanical state of this organism. Here, we perform a mechanical characterization using a combination of microaspiration setups and numerical simulations. We finally propose a description of what happens at the mechanical level during *Hydra* regeneration, allowing quantitative approaches questioning the role of mechanical cues in axis definition.

## Introduction

*Hydra vulgaris* has long been a model of choice in developmental biology because of its remarkable regenerative capabilities (1, 2). Almost any excised tissue piece as well as cellular re-aggregates are capable of reforming a fully viable adult in just a few days. In the former example, an excised tissue piece folds back into a closed spherical shape with both its epithelial monolayers, endoderm and ectoderm, engulfing a water-filled lumen. In the latter, the same tissues start by spontaneously sorting into their relative positions (3, 4) before expelling excess cells to reform a hollow sphere. At this point, both regenerative trajectories converge and these tissue spheres start undergoing osmotically-driven oscillations during which they swell because of water flowing from the environment to the lumen up to the point where the tension building up within the tissues becomes too large and leads to a local rupture (5, 6). After rupture, the samples deflate, close the resulting wound and start another swelling-rupture cycle. These osmotic oscillations induce deformations and therefore stresses within the tissues with relative changes in the radii of the spheres up to 30%. These have been referred to as phase I oscillations and are characterized by a high amplitude and a low frequency (6). Indeed, after a few of these cycles, the oscillations clearly change and enter phase II where they have lower amplitude and higher frequency (6).

In parallel to these mechanical oscillations, the samples establish a chemical pattern involving some characterized morphogens to define the oral-aboral axis of the organism. Most notably, a local expression of HyWnt3 has been shown to be an early signal of axial patterning with this activation defining the future position of the head organizer (7). This local activation is then followed by the establishment of various chemical gradients within the spheres effectively patterning the whole axis (8, 9). At that point, the originally symmetrical samples start elongating in an oblong shape at which ends the adult organs of the head and foot will be regenerated, and the patterning can be considered complete. The switch between phase I and phase II oscillations was thought to be a signature of the establishment of axial patterning (6, 10). Since small excised tissue pieces showed both phases, it was thought that they underwent spontaneous symmetry breaking and retained no memory of axial patterning just as cellular re-aggregates (5, 6).

The question was then to determine how the spherical symmetry was broken during phase I oscillations and how the local head organizer was defined. For a long time, the main hypothesis was that of a purely biochemical spontaneous symmetry breaking in the form of Turing-instabilities of an unknown reaction-diffusion system. In his seminal work, partially inspired by *Hydra* regeneration, Turing has shown how a system of two interacting and diffusing chemical species, which he named morphogens, could become unstable in their homogenous state and spontaneously start forming structures such as dots or stripes (11). These ideas were critical in developing the field of pattern formation and have been adapted to a wide variety of systems and organisms (see for example (12, 13) for recent reviews in the context of developmental biology). In this historical development, *Hydra* has retained its place as a model organism. Notably, Gierer and Meinhardt have expanded on the seminal ideas of Turing and developed a modified version of his reaction-diffusion system specifically designed to explain regeneration and grafting experiments in *Hydra* (14). As a result, the so-called Gierer-Meinhardt model has long been the gold standard in the field. At its inception, it was purely speculative but, since then, some proteins have been shown to possess many of the characteristics required by the Gierer-Meinhardt model. Most notably, the protein HyWnt3, involved in the canonical Wnt pathway is now generally considered to be the activator represented in the Gierer-Meinhardt model (15) since its expression is restricted to the head organizer, it has self-activating capacities and it is the first temporal signature of symmetry breaking during regeneration. However, the necessary long-range inhibitor is notoriously missing. Some promising candidates were put forward, such as Dickkopf (Dkk) (16), Sp5 (17) or *Hydra* astacin-7 (18) but none of these could reproduce the predictions of these models, mostly because of differences in expression patterns.

These models also ignore the mechanical aspects of the process, most notably the osmotic oscillations although it is now well established that they are necessary for proper regeneration (19, 20). This observation suggested a possible coupling between the mechanical state of the tissues and their biological response, as has been observed in a variety of contexts and organisms with impacts on cell division or gene expression including in the canonical Wnt pathway (21–24). Recent results have demonstrated such a coupling in *Hydra* by which the expression of HyWnt3 is reduced when osmotic oscillations are blocked (20) providing a potential direct coupling between tissue mechanics and chemical patterning.

Recent modelling efforts have thus been made in order to incorporate mechano-chemical couplings (19, 25, 26), for instance by making the diffusion constant of the morphogens within the tissues dependent on tissue stretch (19). In most cases though, these couplings were not grounded in experimental evidence. In addition, recent experimental results have started to question the assumptions described so far and underlying these models. First, it was shown that the shift from phase I to phase II oscillations was not a direct signature of axial patterning being established. Instead, the onset of phase II is due to the early apparition of the *Hydra* mouth (27) allowing it to regulate its osmotic imbalance by mouth opening rather than by tissue rupture. One consequence was that axial patterning had to be anterior to the shift between oscillation phases. It then became unclear whether small, excised tissue pieces really went through *de novo* patterning or if they could inherit this patterning from their original host organism. It now appears that they do retain axial patterning although the exact mechanism by which they do so remains unclear, whether by the organization of their ectodermal actin structures called myonemes (28) or by pre-existing biochemical gradients (29).

Although small excised tissue pieces retain axial patterning, it still remains that 1-cellular re-aggregates which cannot conserve either supracellular actin structures or chemical gradients have to show *de novo* axial patterning during osmotic oscillations, 2-there is evidence of a direct coupling between tissue deformations and Hywnt3 expression (20), one of the most important morphogens involved in *Hydra* patterning and 3-osmotic oscillations are required for the proper elongation, morphogenesis and regeneration of both excised tissue pieces and cellular re-aggregates. For all these reasons, the focus of the field is currently shifting to an integrated view of *Hydra* regeneration as a mechano-biological process (26, 30). One clear roadblock to the development of these ideas is our lack of understanding of the rheological properties and mechanical state of regenerating *Hydra* tissue spheres. This often leads to assumptions as to the rheology of these samples and rough estimates of their key mechanical parameters. One exception is the development by Veschgini and colleagues of a two-fingered robotic hand allowing them to apply known, constant deformations on the tissue spheres and measure the resulting forces (31) giving the first local, quantitative measurements of mechanical features in *Hydra*. These measurements were not, however, used to deepen our understanding of the spontaneous osmotic oscillations.

In this work, we try to overcome these limitations and to offer a quantitative characterization of the mechanics of *Hydra* tissue spheres. To do so, we used the well-established micro-aspiration technique (32–35) which we adapted to increase its throughput through the use of original microfluidic constructs following (36). We found three different mechanical behaviors as the aspiration pressure was increased: first an elastic response, then a viscoelastic one and finally tissue rupture, as observed in phase I oscillations. Combining our experimental observations and measurements with the development of a rheological model of elastic shells and numerical simulations, we obtained quantitative measurements of both the main rheological parameters of *Hydra* tissue spheres and the critical pressures required to switch between the three regimes. Thanks to these results, we provide a description of internal tissue mechanics, strains and stresses during osmotic oscillations and reveal that the tissue spheres behave largely as hyperelastic spherical shells during these oscillations. Hopefully, this mechanical characterization will serve as a stepping stone for the study of mechano-biochemical couplings in a quantitative manner.

## Material and Methods

### Hydra maintenance and lines used

3 *Hydra vulgaris* lines were maintained and used for experiments: a watermelon (WM) line (ectoderm GFP / endoderm DsRed2), a reverse watermelon (RWM) one (ectoderm DsRed2 / endoderm GFP) and the AEP line from which embryos were obtained for making transgenic animals. *Hydras* were kept in *Hydra* medium (HM), which consists of 1mM CaCl2, 0.1mM MgCl2, 0.3mM KNO3, 0.5mM NaHCO3 and 0.08mM MgSO4 at a PH between 7 and 7.3. Cultures were stored at 18°C in the dark in an incubator (Pol-Ekko-Aparatura). Animals were fed two to three times per week with newly hatched Artemia (Hobby) and cleaned every two days by changing the medium in which they sit. Animals were starved for at least 24h before the start of any experiment.

### Preparation of tissue spheres

Tissue spheres were made by cutting a whole animal using a sterile scalpel (Holtex, Bistouri UU n°10), as follows: The head and tail were removed by two transverse cuts and the remaining body column was first sliced into 3 or 4 pieces. These pieces were cut further to obtain 6 to 8 samples per adult animal. These samples were left to fold into tissue spheres for 3 to 4 hours in HM.

### Design and fabrication of the removable inserts

Sliding elements containing the cylindrical pipettes were fabricated in dry film according to the protocol detailed in (36). Briefly, photolithography was realized on a 500 μm-thick photosensitive dry film (DF series), laminated on a silicon wafer. Lateral dimension of the photolithography mask defined the length of the insert (20 mm) and its height (1 mm), as well as pipette diameter (50 and 100 μm were used). 100 μm-wide square openings placed at the bottom of the insert were also used for imaging purposes in Movie S3 and Fig S4. After post-exposure bake on a hot-plate, the inserts were released from the wafer during overnight development in an acetone bath that both revealed the patterns and unstick the dry films from the wafer. A single fabrication run allowed to fabricate 100 such sliding elements.

### Mounting and preparing of the microfluidic device

We used a microfluidic tool that allows several micro aspirations to be performed in parallel. This system is composed of a chamber crossed by a removable insert bearing multiple tunnels allowing suction to happen. Both the chambers and inserts were designed using the CAD software Inventor 2020 (Autodesk). The chamber is composed of a main channel which is 20.5mm long, 4mm wide and 0.5mm high (see Fig S1 for the schematics). Halfway through its length, it is barred by another channel which will host the removable insert. The perpendicular channel is 13mm long, 0.5mm wide and 1mm high (Fig S1).

From this design, a mold was manufactured by micromilling using the CNC Mini-Mill/GX (Minitech machinery corporation) with a two size cutter of 1mm diameter on a brass template. The chamber was then made of polydimethylsiloxane (PDMS) (Sylgard 184, Dow Corning) supplemented with 10% curing agent (Dow Corning) from the mold. The PDMS was left to solidify overnight at 70°C and the channel was manually peeled. Inlets and outlets were created by manual punching using a 1mm wide puncher (Harris Uni-Core) at each end of the main channel. The insert channel was opened by manual, transverse cutting to allow insertion of the sliding element. The resulting channel was then bound to a thin PDMS layer, itself bound to a glass slide (Biosigma VBS655/A) by using an oxygen plasma cleaner (Harrick plasma PDC002 with the associated PlasmaFlo gaz mixer). The thin PDMS layer was used to facilitate the introduction of the insert by avoiding direct friction with the glass slide and to avoid leakage. This introduction is done manually with tweezers and is aided by the fact that the insert cannot deviate from its associated channel because of the height difference between both channels. The outlet was connected to a plastic tube of 0.5mm internal diameter and 1.25mm external diameter, sealed at the other end by a 0.6x30mm needle (BD, microlance 3) attached to a 3mL syringe (Braun, Omnifix3P). The needle is deliberately slightly wider than the tube to ensure a hermetic connection. The whole setup was placed in a 100mm Petri dish and immersed in HM. By slowly removing the syringe’s plunger, the channel started filling with HM from the inlet all the way to the syringe while avoiding to create air bubbles in the main channel and checking for proper sealing of the system. Once the plunger was fully removed, the height difference Δ*H* between the water in the syringe and the water in the Petri dish (see Fig S2 for a schematic of the whole setup) was manually controlled and created a pressure difference between chamber inlet and outlet according to:

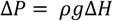

Where *ρ* is the HM density and *g* the gravitational acceleration.

### Micro-aspiration experiments and data analysis

The syringe was first placed 4cm below the water level of the dish to create a light water flow induced by hydrostatic pressure. Several *Hydra* tissue spheres were manually pipetted into the inlet and were thus naturally driven to the holes in the inserts by the water flow, at which point the height difference was set back to 0 for a couple of minutes. A typical experiment could then be started by the application of a controlled pressure by moving the syringe down again. Images were captured under a microscope (Zeiss Axio Vert.A1 equipped with an Axiocam 202 camera). In most experiments, timelapses were recorded in fluorescent microscopy using WM and RWM lines.

In the case of elastic measurements, increasing steps of pressure were applied onto the same samples. Each step lasted for 10min and images were recorded every 10s to make sure that no flow was observed on the samples. After these 10min, the samples were released for 5min to allow them to regain their original shape before applying a new, larger pressure difference.

For visco-elastic measurements, a single pressure step was applied and samples were imaged by fluorescent microscopy for 30min with a 10s time step and the dynamics of the aspirated length was monitored.

To automate the measurement of the aspirated length in these experiments, kymographs of the aspirated tongue were generated in ImageJ (NIH), binarized and analyzed in Python with custom made code to extract the length of the tongue in each image. For elastic measurements, the average length over the last 5min of recording was used whereas for visco-elastic measurements, the whole dynamics was kept.

The acquired images were also used to measure the mean projected area *A* of each analyzed sample by usual thresholding techniques in ImageJ. This measurement was then turned into an average radius

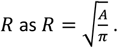

### Osmotic oscillations measurements

To measure osmotic oscillations shown in Fig S8, *Hydra* tissue spheres were prepared as described above. In parallel, a 1% agarose gel was prepared at the bottom of a Petri dish, left to solidify and was manually punched to create 1mm wide wells which then hosted the samples. These wells were used to limit the samples movement during osmotic oscillations. They were then imaged for 24h at a 15min interval. The resulting images were thresholded and analyzed in ImageJ to extract the projected area as a function of time *A*(*t*). Finally, the samples’ radii *R*(*t*) were computed as described above. The equilibrium radius *R*_0_ of a given sample was defined as the minimum of *R*(*t*) over the whole recording allowing us to estimate the strains on the inflating sphere.

### Finite element simulations

Numerical simulations were performed with the finite-element solver Comsol Multiphysics (COMSOL, Inc). The micropipette was represented as a rigid, motionless cylinder of inner radius *R*_*p*_= 50*μm*. The reference state of the *Hydra* sphere was a shell of outer radius 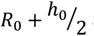 and inner radius 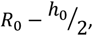, whose center *O* was located on the cylinder’s longitudinal symmetry axis *O*z. We used contact boundary conditions ensuring that the (deformed) *Hydra* sphere and the micropipette never overlap. *Hydra* tissue was modeled as a hyperelastic Saint-Venant Kirchhoff material. We denote by *E* and *v* the Young’s modulus and Poisson’s ratio of the tissue and considered nearly incompressible conditions (*v* = 0.495). The external pressure Δ*P* was applied on the spherical section contained within the micropipette. Simulations were performed in 2D, enforcing the rotational invariance of the system (*Hydra* + micropipette) about *O*z. The equations of elasticity were treated within a pressure formulation, adapted to approximately incompressible elastic materials. For each value of the applied pressure, we recorded the maximal deformation observed on the symmetry axis denoted *δ*z(*r* = 0), and computed as a spatial average along the shell’s thickness. We checked that our results were robust to smoothing the sharp corner of the micropipette by a fillet over a scale of the order of 5*μm*. They were also insensitive to decreasing the typical mesh size.

### Extraction of rheological parameters

To measure Young’s moduli in the elastic phase, each sample was treated separately. Based on the acquired images, we measured their projected area *S*_0_and turned them into rest radii 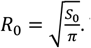 . Then, we used our measurements of Δ*P* as a function of *δ*z (Eq 7) and fitted them by a straight line in Python. The slope of this straight line was kept and multiplied, according to Eq 7, by 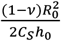 to extract the Young’s modulus of that specific sample. We then repeated the procedure on many samples and report the mean and standard deviation of this distribution in the text.

In the visco-elastic phase, we measured the dynamics of the aspirated length as a function of time *L*(*t*) and fitted them in Python according to Eq 8. To extract Young’s moduli, we used the same formula as above except that we replaced the slope of a linear fit of several points by the ratio of Δ*P* over *δ*(1 − *κ*), *δ* and *κ* stemming from the fit of the full dynamics.

To estimate the order of magnitude of the effective viscosities in the same experiments, we used the same fit according to Eq 8 to extract the speed of tissue flow *U* and estimated the viscosity as 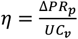 with *C*_*v*_= 1.

### Measurement and data analysis of threshold pressure measurements

For the measurement of the pressure thresholds distinguishing each regime, we used the fraction of pieces changing their rheological behavior at different applied pressures. To observe the switch between the elastic and viscous regimes the pressure was gradually increased by lowering the syringe in 3cm increments, starting low enough that no viscoelastic behavior was observed (usually 10 or 14cm). At each pressure level and after 20 minutes, we noted the fraction of pieces that had changed regime, and the experiment was stopped when all samples had done so. Kymographs were used to visually separate both regimes. The length profile over time was constant in the elastic regime, whereas it evolved linearly with a clear slope in the viscous regime.

The same strategy was used to measure the rupture threshold, but we started at higher applied pressures. Images were taken in transmission, as it facilitated the observation of briefly detached pieces of tissue in the holes of the inserts, the first sign of the rupture regime. They were also taken this time at a rate of ten minutes of suction per threshold. Identifying the switch between the viscous and the rupture regime was done visually.

In both cases, we thus ended up with a fraction of samples adopting a certain behavior as a function of applied pressure. Assuming that individual thresholds are variable and normally distributed, this cumulative frequency followed a sigmoid shape of the form 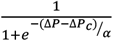 where Δ*P*_*c*_is the threshold pressure at which 50% of samples have switched behavior and *α* is a parameter controlling the steepness of the sigmoid function. We thus fitted our data with that functional form using Matlab’s (Mathworks) curve fitting toolbox. In the manuscript, we report threshold pressures as Δ*P*_*c*_resulting from the fits and error bars represent the 95% confidence interval of these fits.

### EDTA/blebbistatin experiments

Rheological measurements on *Hydra* whose cell adhesion had been inhibited were carried out by immersing the tissue spheres in solutions of either 2 mM EDTA (Sigma-Aldrich) or 5μm blebbistatin (Tocris Bioscience) in HM. The entire microfluidic system was immersed in the solution, and the *Hydra* pieces were left to bathe in it for around ten minutes before aspiration to give the drug time to take effect before measuring. The rest of the protocol was identical to that of the rheological measurements explained above.

### Spinning disk imaging

To estimate the thickness of *Hydra* tissue spheres, several WM samples were imaged underneath an Eclipse Ti2 microscope (Nikon) equipped with a CSU-W1 spinning disk unit (Yokogawa) and an Orca Fusion BT camera (Hamamatsu). Z-stacks were automatically acquired with a step of 1 micron and vertical slices were made using ImageJ (Fig S4). The thickness of six samples were manually measured using the same software and were all found to be in the 15 − 25*μm* range.

### Mechanical description of *Hydra*tissue spheres

We consider a thin spherical shell of undeformed inner and outer radii 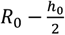 and 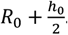. Normal components of the Cauchy stress tensor are denoted *σ*_*a*_(*a* = 1,2,3). Briefly, we re-express the strain energy function, Eq 4, of a Saint-Venant Kirchhoff material as a function of the stretch ratios 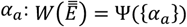. Denoting *P* the shell pressure field, stress components are given by 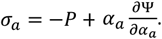 . By the assumption of plane stress, *σ*_3_ = 0, we obtain the expression of *P* and deduce the circumferential, or hoop stress *σ* = *σ*_1_ = *σ*_2_. For an incompressible material, 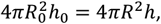, where *R* and *h* denote the deformed radius and thickness, respectively. We note 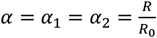 the circumferential stretch ratios, and deduce the normal stretch ratio 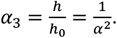. The calculation follows (37, 38) and yields the expression of the hoop stress of a Saint-Venant Kirchhoff spherical shell, as given by Eq 11.

## Results

### Multiplexed micro-aspiration setup

In order to define and measure the rheological properties of *Hydra* tissue spheres, we settled on the use of micro-aspiration experiments. These experiments are now well-established, robust and measure rheological properties at length scales and frequencies relevant for the regenerative process. In contrast with Atomic Force Microscopy (AFM), previously used to locally probe the stiffness of adult *Hydras* (39), our experiments were particularly designed to study the mechanics of tissue spheres during the osmotic oscillations which is why we employed elongations rather than indentations and probed the large deformation regime. Their main drawback is their intrinsically low throughput. In their original form, micro-aspiration experiments can only probe one sample at a time with a single experiment running for around 1h (4, 33). We thus started by developing and adapting a microfluidics device making use of removable inserts (36) to parallelize the experiment, in the same spirit as the technique developed in (40) but with circular pipettes which prevent singularities and leakage. This device was composed of two different objects. First, a microfluidic channel was designed to host the samples, it was 3cm in length and 500μm in height (Fig 1A, Fig S1). Halfway through this channel and perpendicular to it sat another channel meant to host the removable insert. This channel was 500μm wide and 1mm high. This whole construct was then manufactured in polydimethylsiloxane (PDMS).

**Fig 1.**
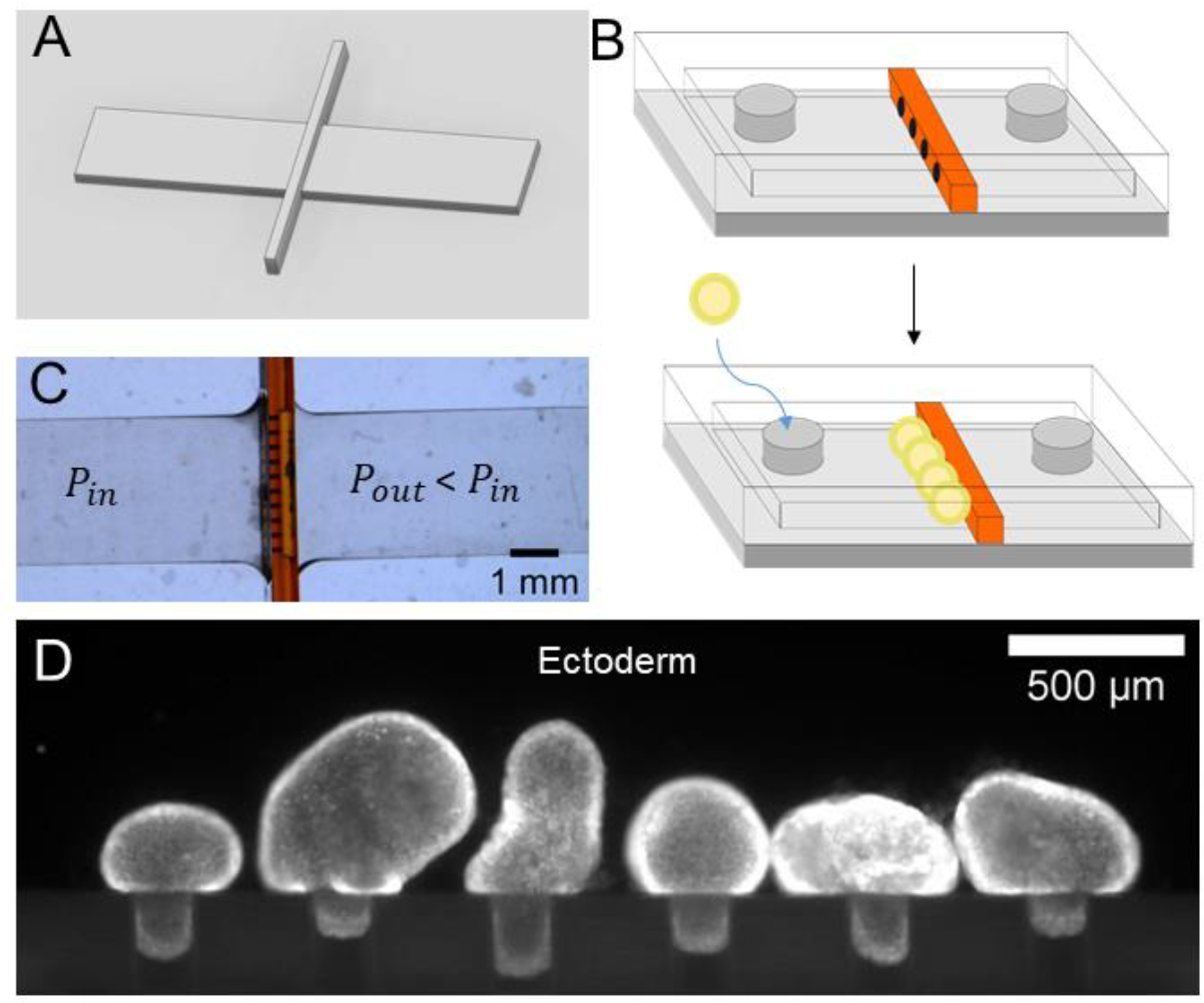
Parallelized micro-aspiration experiments. A: Schematics of the mold for the PDMS channel (details on dimensions can be found in Fig S1). B: Representation of the principle of the experiment, the insert, in orange, effectively creates an array of cylindrical tunnels, equivalent to model micropipets. The tissue spheres, in yellow, flow towards these openings and, once sealed, get aspirated within. C: Snapshot of a mounted channel with an insert containing ten tunnels in parallel. D: Fluorescent imaging of the ectoderm of six samples aspirated within the tunnels.

Most of the experiments presented here were carried out with removable inserts, microfabricated in dry film, bearing either 6 or 10 circular openings, 500μm or 300μm away from one another, 300μm in length and 100μm in diameter. Of note, these holes sat, vertically, at the midpoint of the samples’ channel (250μm, Fig 1B). Tissue spheres aspirated in these holes therefore didn’t touch either the bottom or the top of the PDMS channel which had no influence on the aspiration.

Using hydrostatic pressure, we applied a difference in pressure between the inlet and outlet of the main chamber (with *P*_*in*_> *P*_*out*_, see Fig S2) which initially created a flow of water from the Petri dish to the outlet of the chamber. Previously prepared *Hydra* tissue spheres were then manually pipetted into the entrance of the main channel. Thanks to the flow of water, the samples naturally aligned with the openings in the insert. Once all openings were blocked by a sample, the flow of water stopped and aspiration of the tissue spheres was observed (Fig 1D, Movie S1). Of note, in most experiments, the pressure difference was set to 0 as soon as the samples aligned with the holes so that a controlled step in pressure could be applied thereafter.

### Rheological behavior depends on applied stress

Using this multiplexed setup, we started by studying the response of newly formed *Hydra* tissue spheres to micro-aspirations in the kPa range using circular openings with a radius *R*_*p*_= 50*μm*. We observed two different rheological responses depending on the applied pressure. Up to around 2.5kPa, tissue spheres exhibited an elastic response in the sense that for a given applied pressure difference, a tongue of tissue was aspirated within the holes which length increased in the first tens of seconds but then saturated and remained constant over much longer time scales. When the pressure was released, we also observed that the samples retrieved their original shape, and that aspiration and release dynamics were very similar (Fig S3), consistent with the reversible behavior of an elastic material. To further make sure that we were not missing a viscous behavior at longer time scales, we managed to trap tissue spheres for tens of hours in this elastic phase without any indication of tissue flow (Movie S2).

However, when we increased the applied pressure, we started observing samples flowing inside the holes, a signature of a viscous behavior. This demonstrates a real change in mechanical behavior of the tissue spheres akin to the behavior of yield stress fluids and requiring two different experimental and theoretical approaches.

Finally, at even higher applied stresses, we started observing rupture of the tissue and cell detachments, reminiscent of the rupture observed under normal phase I osmotic oscillations (Movie S3). Since all three behaviors seemed relevant to understand the mechanical behavior of normally regenerating tissue spheres, we decided to characterize them all, as well as the pressure thresholds between these different phases.

### Elastic behavior at low applied stresses

To characterize the elastic behavior of the tissue spheres, we applied steps of pressure on the same samples and recorded, for each step and for each sample in an experiment, the length of the aspirated tongue *L* (Fig 2A). We monitored the samples closely to make sure they did not start flowing in the holes, in which case their analysis was stopped. As expected, we found that the higher the pressure, the longer the aspirated tongue (Fig 2A). To estimate a Young’s modulus from these measurements, we needed a rheological model of an elastic spherical shell of mean radius *R*_0_ and thickness *h*_0_.

**Fig 2.**
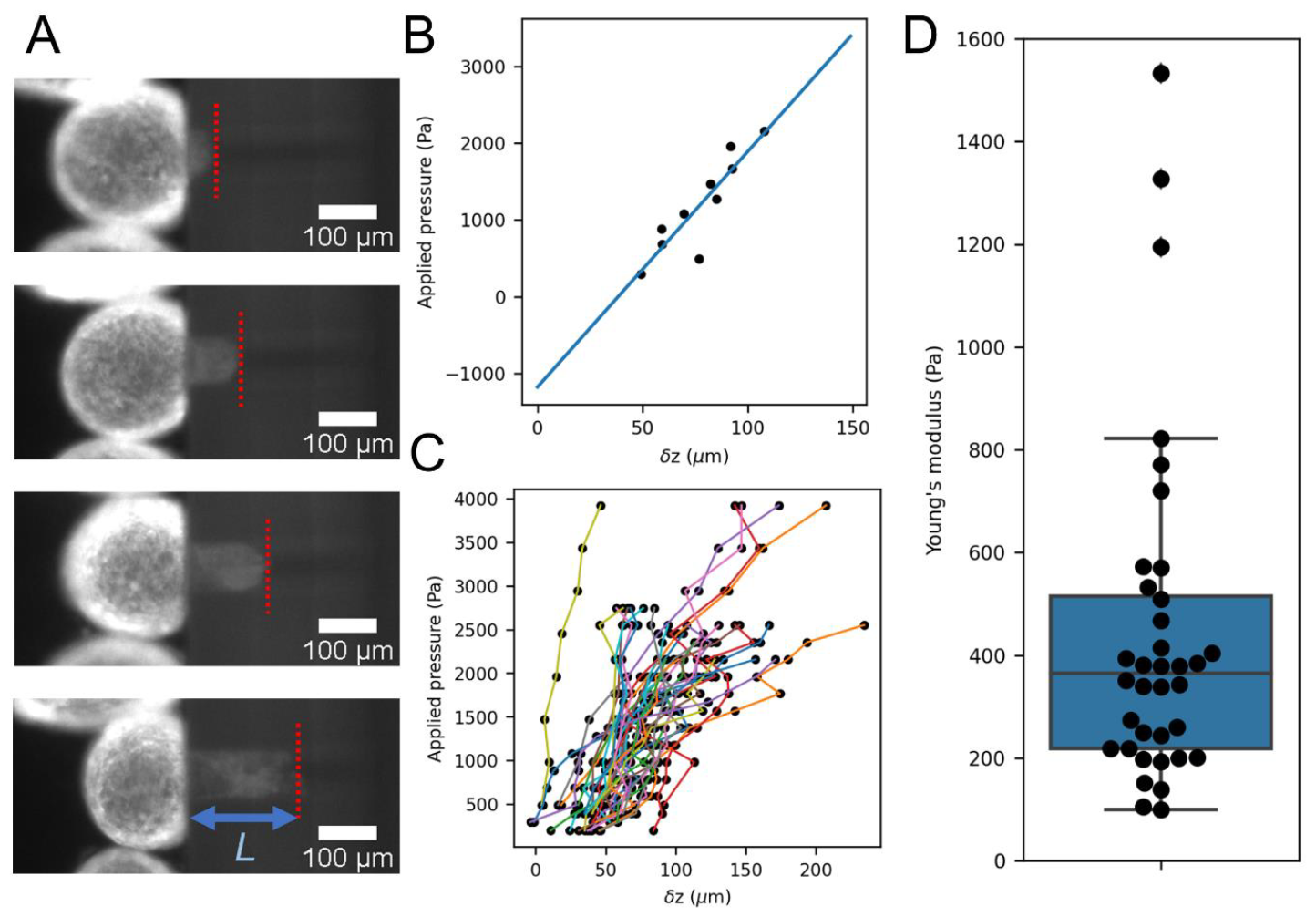
Elastic behavior quantification. A: snapshots of one tissue sphere aspirated in a hole at increasing pressure steps (top to bottom). The dashed red line shows the position of the aspirated tongue at each step and L is the aspirated length. B: Linear relationship between the applied pressure and δz for a single tissue sphere and 10 different pressure steps showing both the linear behavior (fit as a blue line) and negative intercept. C: Same measurement as in B, repeated on n=36 tissue spheres, each represented by a differently colored line. D: distribution of Young’s moduli obtained by this method. Black dots are individual measurements, the box plot shows the median and quartiles of the distribution. We found E= (4.4 ± 3.3).10^2^ Pa (mean ± standard deviation, n=36).

As is the case for cell aggregates, we expect that intercellular adhesion and cellular cortical tension will contribute to an effective surface tension of the *Hydra* tissue spheres, which we denote by *γ*.

A pressure difference Δ*P* through the micropipette of radius *R*_*p*_generates an aspiration force 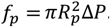. For an aspirated length *L*, the free energy *F* of the aspirated shell reads:

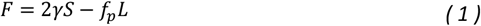

Where *S* denotes the surface area of the sample and the factor 2 takes into account the presence of two interfaces. Assuming incompressibility, the volume *V* of the sample is constant. For small enough deformations during aspiration, we expected that the thickness *h*_0_ of the sample would remain constant. To validate this hypothesis, we performed spinning disk confocal microscopy on normal tissue spheres and aspirated ones to reconstruct their 3d structure and estimate their thicknesses. We found *h*_0_ = 20 ± 5*μm* in both cases and didn’t observe any clear change due to micro aspiration (Fig S4). Since *V* = *Sh*_0_, the surface area of the sample also remains constant *dS* = 0. In other words, in the case of the small deformations of an incompressible shell of approximately constant thickness, tissue surface tension does not contribute to the total force *f* exerted on the sample:

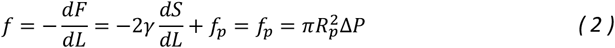

Indeed, our experiments do not exhibit a clear half-sphere of radius *R*_*p*_, typical of the early entry of a full cell aggregate into a micropipette (41). In addition, (33) elegantly demonstrates that surface tension manifests itself as a critical pressure below which no aspiration is observed and which needs to correct all subsequent applied pressures. The fact that we observed aspirations with applied pressures as low as 200Pa also indicates that surface tension can largely be ignored in our experiments.

The rest state of the system is simply that of the initial spherical shell of radius *R*_0_ and due to that initial curvature, the displacement of the tip of the tissue tongue inside the micropipette *δ*z reads:

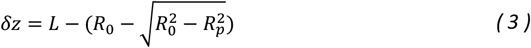

As expected for an elastic material, we found a linear relationship between this displacement and the applied pressure Δ*P* (Fig 2B). We also noticed that for many samples, our linear fits had negative intercepts at the origin. This could be the signature of a stress-stiffening effect by which the effective Young’s modulus would increase with the applied pressure, *i.e.* the existence of non-linear elastic behavior, as observed in suspended cell monolayers (42).

To take into account this observation, we turned to a nonlinear, hyperelastic description of the *Hydra* tissue, using, as previously advocated by (43), the Saint-Venant Kirchhoff model, defined by the strain energy function:

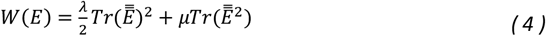

where *λ* and *μ* denote the Lamé coefficients, and 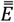 is the Green-Lagrange strain tensor (37). This equation reduces to its linear elastic counterpart for small deformations, when 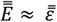 with 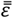 the infinitesimal strain tensor. Of note, the Saint-Venant Kirchhoff model is defined by the same parameters as would be the case for a linear elastic material, with the following classical relationships between Lamé coefficients and other elastic moduli: *E* and *v*, respectively, the Young’s modulus and the Poisson’s ratio of the sample.

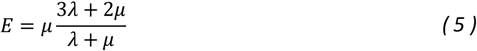

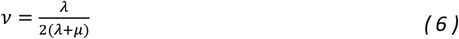

In analogy with the behavior of linearly elastic thin spherical shells (see for instance (44)), and taking into account the non-zero intercept Δ*P*_0_ of the linear fits of our data (Fig 2C, Δ*P*_0_ = (−6.4 ± 1.3). 10^2^ *Pa*), we expect that the displacement and the applied pressure should be related according to:

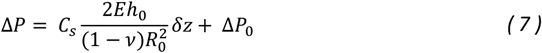

Where *C*_*s*_is a dimensionless, geometry-dependent pre-factor. Of note, in this description, we ignore the complex composite structure of the shell which includes two different epithelial layers as well as the extra-cellular matrix.

In order to estimate *C*_*s*_, we performed numerical simulations of micro-aspiration of hyperelastic spherical shells by a single cylindrical tunnel (Fig 3A). In our experimental data, we had *R*_0_ = 162 ± 28*μm* (mean ± standard deviation, n=36) and *h*_0_ = 20 ± 5*μm*. As we had no measurement of the Poisson’s ratio of our samples, we made an assumption of near incompressibility and set *v* = 0.495. Based on these values, we submitted spherical shells to applied pressure Δ*P* and measured the resulting *δ*z. As shown in Fig 3B, these simulations confirmed the linear relationship between Δ*P* and *δ*z over a similar range of applied pressures and displacements and allowed to estimate *C*_*s*_thanks to the knowledge of the value of *E* and Δ*P*_0_. Of note, other element-based numerical simulations were performed with parameters varying in the following ranges: 25 ≤ Δ*P* ≤ 2000*Pa*, 15 ≤ *h*_0_ ≤ 25*μm*,130 ≤ *R*_0_ ≤ 200*μm*, 200 ≤ *E* ≤ 750 *Pa* and with a fillet size varied between 2 and 10*μm*. Overall, our estimate of the coefficient *C*_*S*_reads *C*_*S*_= 23.4 ± 4.3, in the vicinity of 23, suggesting that the order of magnitude of our results was unaffected by uncertainties on these different parameters. Further, our estimate of the intercept in these simulations is Δ*P*_0_ = (−8.5 ± 2.1). 10^2^ *Pa*, consistent with the above experimental values.

**Fig 3.**
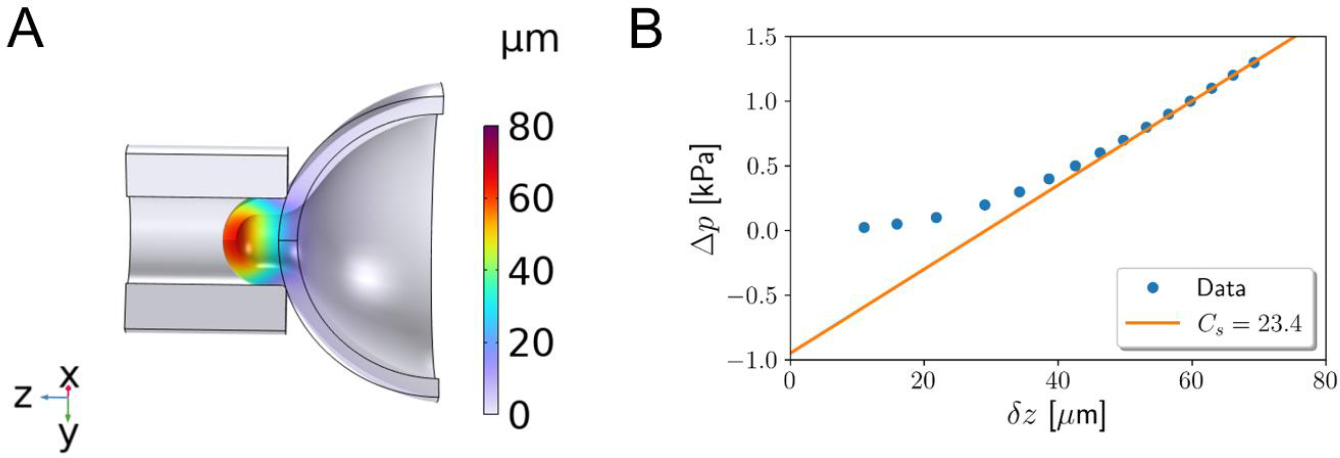
Numerical simulations. A: Graphical representation of the deformed shell under. The color code represents the z-component of the displacement field. Black lines delineate the reference state of the system. Parameter values for these simulations are R_0_ = 160 μm, *h*_0_ = 20 μm, E = 440 Pa, v = 0.495, ΔP = 10^3^Pa. B: applied pressure Δp versus the displacement δz for the same parameter values. The slope of the fitted line, in orange, corresponds to a value C_S_ = 23.4.

Thanks to our multiplexed setup, we were able to measure up to six samples in parallel in one sitting. By then repeating these experiments eight different times, we achieved a total of 36 tissue spheres (Fig 2C) characterized in that phase. For each of them, we extracted the slope of the linear relationship between Δ*P* and *δ*z, its initial radius *R*_0_ and finally obtained a quantitative measurement of its Young’s modulus using Eq 7 (Fig 2D). From the above estimate of the coefficient *C*_*S*_, we found the Young’s modulus of *Hydra* tissue spheres to be (4.4 ± 3.3).10^2^ Pa (mean ± standard deviation, n=36).

We also observed a large variability in our measurements of individual Young’s moduli (Fig 2D) which could hint at subtle effects missed by our averaging approach. The different tissue spheres could differ in initial size or degree of inflation since we do not precisely control the time they need to fold back into tissue spheres after being cut. To better understand this variability, we looked at the correlation between the samples’ radii and their extracted elastic moduli. We indeed found a positive correlation, albeit weak, between these measurements (Fig S5). We then asked whether this correlation could be due to differences in inflation state. To do so, we ran new experiments measuring the elastic moduli of tissue spheres whose oscillations were blocked by the addition of 70mM sucrose in the outside medium. Doing so, we found 1-that the obtained elastic moduli distributions in both cases were not statistically different (Fig S6) and 2-that addition of sucrose abolished neither the weak positive correlation between size and elasticity (Fig S5) nor the existence of stiffer samples. We conclude that some, but not all, of the observed variability comes from differences in initial size rather than inflation rate between different samples. A better understanding of this dependency would be a natural continuation of this work.

In addition, we validated the relevance and precision of our experimental procedure by repeating elastic measurements under two chemical modulations: EDTA and blebbistatin. EDTA is a chemical well-known for weakening cell-cell adhesions in many epithelial tissues (45) whereas blebbistatin inhibits acto-myosin contractility (46). We expected that both treatments should lower the samples’ elasticity either by allowing larger tissue-scale deformations in the case of EDTA or cell-scale deformations in the case of blebbistatin by making the actin cortex of each cell softer. We indeed found a clear and statistically significant decrease in Young’s modulus with both treatments (Fig S6, EDTA: E=(3.0 ± 2.0).10^2^ Pa, blebbistatin: E=(1.9 ± 0.7).10^2^ Pa), confirming that the observed elastic behavior of the tissue spheres stems from a complex interplay of single cell mechanics and tissue structure.

Overall, we characterized in a quantitative way, the response of *Hydra* tissue spheres to applied stresses and found them to behave hyperelastically, *i.e.* as soft (*E*∼500*Pa*) elastic materials at small deformations, in line with AFM measurements on adult *Hydras* (39), while they become increasingly stiffer as the applied pressure increases.

### Visco-elastic behavior at intermediate applied stresses

We then turned our attention to the next observed behavior which involved flowing of the samples within the tunnels. This visco-elastic behavior has long been observed and studied, both on single cells (32, 47, 48) and multi-cellular spheroids (4, 33, 41). We followed the usual approach in that situation where a single, controlled pressure step was applied in the device at t=0 after the samples were already placed close to the entrance of the tunnels. The dynamics of the aspirated length *L*(*t*) was then recorded by fluorescent microscopy for tens of minutes (Fig 4A). It was important to keep recording on these longer time scales in order to be certain of the behavior adopted by each sample as a very slow flow could easily be misinterpreted as an elastic response if the observation time scale was too short.

**Fig 4.**
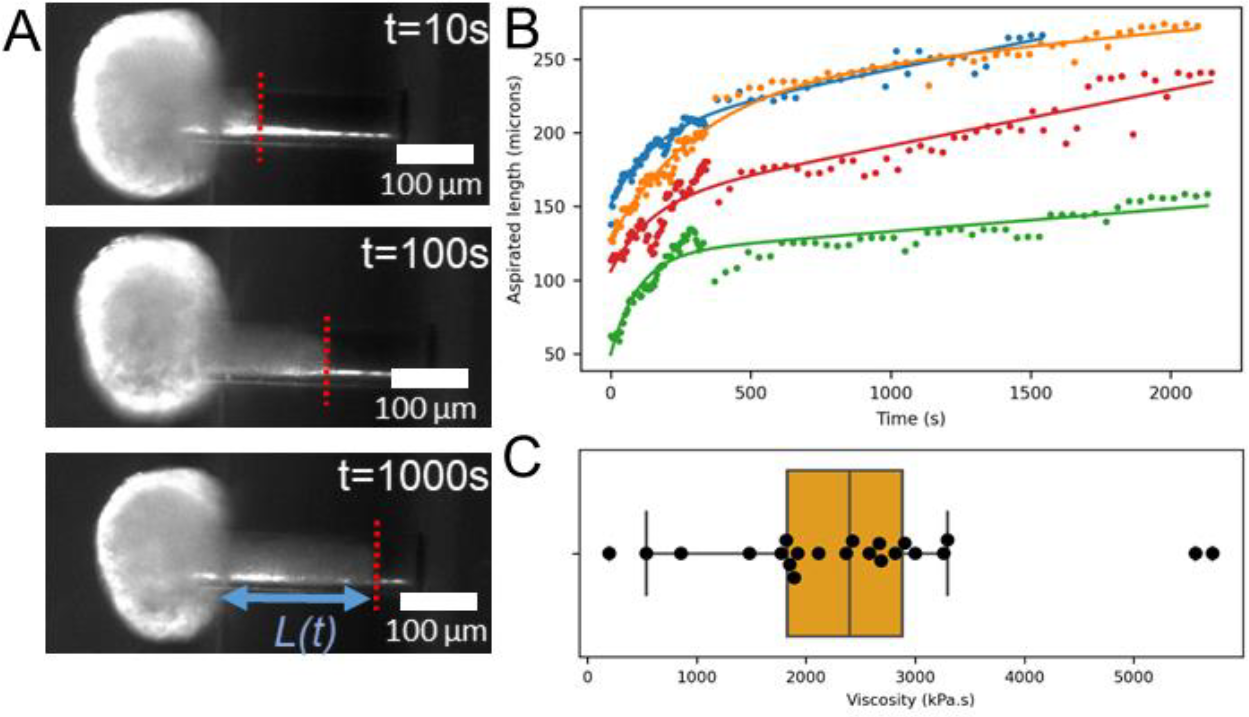
Visco-elastic flow at large applied pressure. A: snapshots of a tissue sphere showing flow at a constant applied pressure. B: four examples of typical dynamics of the flow as a function of time. Dots of different colours represent four independent samples and solid lines fits by Eq 8. C: distribution of measured viscosities as black dots with box-whisker plot in orange. We found it to be (2.4 ± 1.2).10^6^ Pa.s (mean ± standard deviation, n=22).

Since the behavior is qualitatively similar to that observed for cellular aggregates aspirated in cylindrical micropipets, we first applied the well-known modified Maxwell model developed in (33). Briefly, a dashpot representing tissue scale viscosity is combined in series with a standard linear solid unit incorporating cell scale viscoelasticity as well as tissue scale elasticity. We fitted our data with the form of *L*(*t*) predicted by this model:

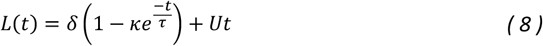

Where *δ* is linked to the short-term elastic deformation, *κ* is a constant linked to the relative contributions of the two springs in the standard linear solid unit, *τ* is the cell scale visco-elastic relaxation-time and *U* is the long term viscous flow rate. We found that this functional form was also relevant to describe our data quantitatively (Fig 4B) despite the morphological differences between our samples (empty shells) and ball-shaped aggregates (full spheres). We could however employ a scaling argument similar to the one applied in this model to estimate an order of magnitude of the effective viscosity of our samples. As for cellular aggregates, friction between the tissue and the tunnels walls can be neglected since the long-term aspirated length scales not as the square-root of time, but as a linear function of time. Assuming that dissipation is due to cell rearrangements at the entrance of the tunnels thus acting on a length scale of order *R*_*p*_, the total dissipative force can therefore be approximated as *f*_*viscous*_≈ *C*_*v*_*R*_*p*_*Uη* with *η* the viscosity of the samples and *C*_*v*_a dimensionless, geometrical pre-factor equal to 3*π* for a ball-shaped aggregate and unknown in our geometry. This viscous force has to be balanced by the aspiration force which we have shown to be 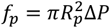 which leads to:

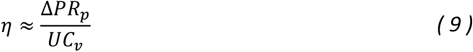

To the best of our knowledge the computation of *C*_*v*_has never been done for spherical viscous shells and in the same geometry as the one in our experiments. We will see below that a precise measurement of this effective viscosity has little impact on the description of *Hydra* osmotic oscillations and we limited ourselves to obtaining an order of magnitude estimation of the viscosity by taking *C*_*v*_= 1. Note that assuming that tissue scale dissipation acts across the thickness h of this multilayered tissue does not change our estimate as h and *R*_*p*_have the same order of magnitude. Again, our microaspiration setup allowed us to efficiently gather data on multiple samples and combining them, we found an estimation of the viscosity of (2.4 ± 1.2).10^6^ Pa.s (mean ± standard deviation, n=22, Fig 4C).

The full dynamics of aspiration in that phase can also yield measurements of the Young’s modulus of the samples. In line with our analysis of the purely elastic response, we expect here that:

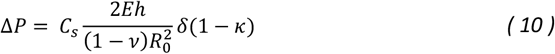

Similar to Eq. 7 with *δ*(1 − *κ*) the immediate elastic aspirated length in the notation of this model. Applying this relationship to the results of our fitted data, we found *E* = (3.7 ± 1.6) . 10^2^*Pa* (Fig S7), in accordance with our results on the purely elastic behavior of *Hydra* tissue spheres.

These visco-elastic experiments were made difficult by the fact that in many instances, we observed, in addition to the flow of the samples, cells detaching at the tip of the aspirated tongue, indicative that the tissues were already rupturing. These cases were obviously discarded from the analysis, limiting the amount of available data. Overall, it appeared that at these applied pressures, the visco-elastic phase and rupture were often occurring together.

### Critical pressures between different behaviors

To further study this question, we then turned our attention to the critical pressures required to transition between the three regimes: elastic, viscoelastic and rupture. We often found cases where, for a given applied pressure, a fraction of the samples loaded in the same setup would behave as elastic solids while other behaved as viscoelastic fluids (Fig 5A). To characterize the first critical pressure, we therefore measured, at different applied pressures, the fraction of samples showing a viscoelastic versus elastic behavior while discarding rupturing ones. We fitted this fraction as a function of the applied pressure by a sigmoid (see Materials and Methods, Fig 5B). We defined the critical pressure as the center of this sigmoid where 50% of samples had switched from one behavior to the other. Following this method, we found the critical pressure between elastic and viscoelastic behaviors to be 2.22 ± 0.14 kPa (error bars represent the 95% confidence interval of the sigmoid fit). We then employed the same strategy to characterize the critical pressure leading to tissue rupture (Fig 5C). We found it to be 2.37 +-0.08 kPa, very close to the previous one confirming our intuition that both behaviors were linked together and that tissue flows inside the holes were already a signature of their lack of integrity.

**Fig 5.**
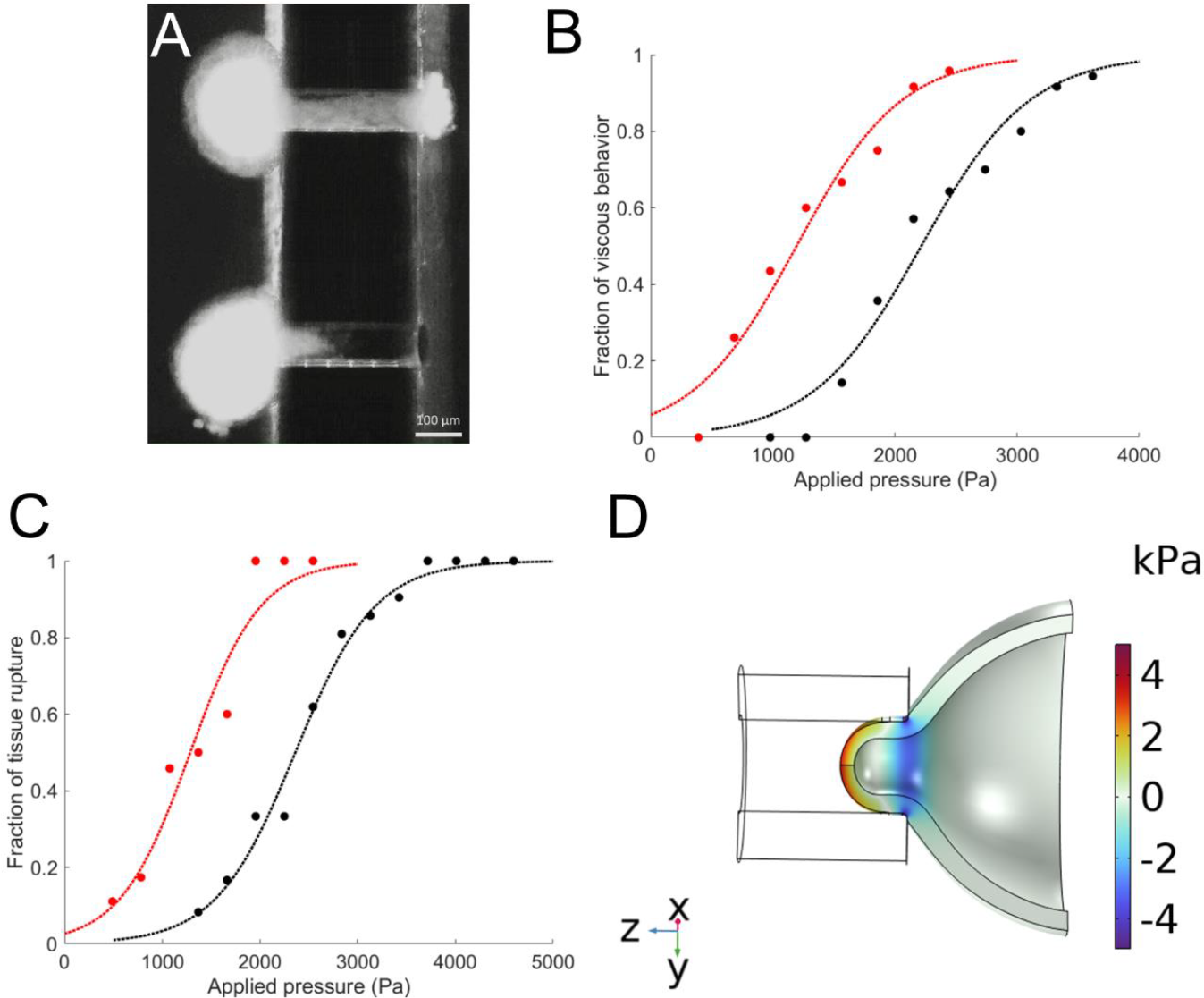
Critical stresses between different mechanical behaviors. A: snapshot of an experiment showing two similar samples displaying different behaviors at the same applied pressure. B: Quantification of the fraction of viscous behavior at different applied pressures. C: quantification of the fraction of samples showing tissue rupture as a function of applied pressure. In B and C, dots are data and solid lines fit by a sigmoid function. In black are control samples and in red samples pretreated with 2mM EDTA. Each point is derived from 18 to 40 samples stemming from 8 different experiments in B and 4 in C. D: Numerical simulations. The color code represents the hoop stress within the aspirated shell when aspirated at a pressure of 1kPa. Parameter values are the same as in Fig. 3.

The most natural explanation for this lack of integrity is the weakening and rupture of cell-cell contacts since we often observed detachment of single cells, making a possible deadhesion of the tissues to the extracellular matrix unlikely. To verify this hypothesis, we repeated our measurement of the critical pressures required to induce either a viscous behavior or tissue rupture on tissue spheres pre-treated with 2mM EDTA as we expected this treatment to facilitate the induction of a viscous behavior and tissue rupture. This is indeed what we found with a shift of the critical pressure from 2.22 kPa to 1.19 ± 0.16 kPa for the viscous behavior (Fig 5B) and from 2.37 kPa to 1.29 ± 0.19 kPa for rupture (Fig 5C).

It is important to note that all these values are difference of pressure applied between the two sides of the holes and not necessarily the stress actually acting in the tissue called the hoop stress. Said differently, the applied pressure required to rupture the tissue shouldn’t be directly taken as a measurement of the ultimate tensile stress of the whole tissue. To relate the applied pressure and internal tissue stresses, we turned again to numerical simulations. Within the range of parameters studied, the maximal values of the hoop stress we found was on the order of a 2-7 kPa (see Fig 5D for an example).

Since in normal conditions, the threshold for viscous behavior cannot be clearly distinguished from the threshold for rupture, our measurements revealed that up to rupture, which is also observed in normal osmotic oscillations, regenerating *Hydra* tissue spheres behave as hyperelastic spherical shells with a stress stiffening effect.

### Mechanical description of oscillating Hydra tissue spheres during regeneration

After folding back into a spherical shape, regenerating *Hydra* tissue spheres experience a series of swelling-rupture cycles which have been shown to be driven by the difference in osmolarity between the inside and outside of the spheres. These oscillations are required for the proper morphogenesis and regeneration of the samples although their exact role remains unknown. Thanks to previous work on the topic (5, 49) and our own mechanical characterization, we can provide a good understanding of the internal tissue mechanics occurring during this phase.

The key quantity is the circumferential or hoop stress, noted here σ, which is known to control tissue integrity and to be able to alter key biological processes. It is given for an incompressible, thin elastic spherical shell of a Saint-Venant Kirchhoff material (see (37, 38) and the Methods section) by the expression:

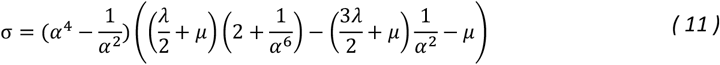

With 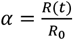 is the stretch ratio, and *λ* and *μ* the Lamé coefficients. This gives the nonlinear constitutive relationship between stresses and strains in the spherical shell during the oscillations which allows to directly extract the hoop stress from a measurement of the radius of the sample as a function of time (see Fig S8). One can check that Eq 11 reduces to the relationship expected of a linear elastic thin shell in the limit of small deformations where |*R* − *R*_0_| ≪ *R*_0_:

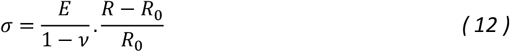

At larger deformations, the tissues get stiffer and the experimentally measured values of *E* imply that, in this regime, the differential elastic modulus 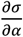 is on the order of 10kPa, in accordance with previous estimates of the sphere’s stiffness during osmotic oscillations (5).

Based on our experimental observations, we will approximate our samples as hyperelastic, symmetric, spherical shells of time-dependent thickness, *h*(*t*) and radius *R*(*t*) (Fig S8) submitted to a difference in osmolite concentrations (*C*_*in*_− *C*_*out*_) > 0. The fact that the spheres are constantly swelling indicates a constant influx of water into their lumen and that the system never reaches osmotic equilibrium. Furthermore, Kücken et al. (5) showed that the increase of *R*(*t*) can be described by a Darcy-type law of the form:

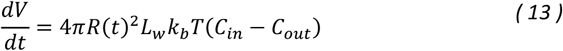

Where *V*(*t*) is the volume of the sphere, *L*_*w*_is the water permeability coefficient, *k*_*b*_is Boltzmann’s constant. The dynamics of swelling is thus entirely controlled by the influx of water which imposes a constantly increasing deformation on the tissue sphere with a growth rate of the radius on the order of 5*μm*/h. This influx of water effectively inflates the sphere and creates a restoring elastic pressure *P*_*el*_which our mechanical description can also predict. Following the mechanics of spherical thin shells, this pressure can be written as:

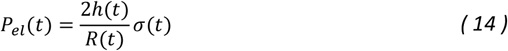

Using the experimental measurement of *E* we previously obtained and the same value of the Poisson’s ratio, *v* = 0.495, we were able to approximate the values of *λ* and *μ* (*μ* = 148*Pa, λ* = 100*μ* ). Using these values and experimental measurements of α(t) during the oscillations allowed us to estimate σ(*t*) and we found it to be on the order of a few kPa (Fig S8). Most notably, this approach allowed us to estimate the critical stress, within the tissue, to induce rupture and we found it to be on the order of 4-6kPa. This is the same order of magnitude as the maximum hoop stress obtained by numerical simulations of the elastic deformation during micropipette aspiration. Overall, we demonstrate that our results allow to map measurements of the radius of the tissue spheres, which are experimentally easy to access, into both internal stresses within the tissue and restoring elastic pressure acting on the fluid inside of the spheres.

## Discussion

The effect of stretching on the biology of epithelial tissues has emerged in recent years as an important topic in developmental biology since developing organisms often undergo substantial deformations which have an increasingly well recognized impact on their biological response. In the context of *Hydra* regeneration, Ferenc et al. observed that the levels of expression of Wnt ligands, especially HyWnt3, are directly linked to the cumulative amount of stretching in the tissue (20). In a similar fashion, stretching adhering monolayers of epithelial cells has been shown to induce cell division through the activation of Piezo1 mechanosensitive ion channels (22) to restore proper cell density. Anisotropic deformations also have a large impact on the tissue as uniaxial cyclic stretch leads to re-orientation of the cells perpendicular to the direction of application (50, 51), as was observed for the mitotic spindle of single cells (52). In many of these experiments, however, stresses within the tissues couldn’t be recorded after stretching.

One notable exception was achieved on freely suspended epithelia axially stretched with a measurable force (42). This experimental setup allowed the authors to characterize the rheology of Madine–Darby Canine Kidney (MDCK) monolayers. Interestingly, they observed a non-linear response with three distinct phases: a linear one at small deformations with a small equivalent elastic modulus, a second linear one for intermediate values of stresses and strains with much higher stiffness and a third one corresponding to the rupture of the tissue. This behavior is very similar to the one we observed in this work. The authors quantified the second phase by linear fitting of the stress-strain curves with negative intercepts, as we did. Since they approximated the tissue in this phase as linearly elastic, they found a high elastic modulus on the order of 20kPa, similar to our observations of *Hydra* tissue spheres differential moduli at intermediate stresses. Finally, the authors found that the monolayer was able to withstand deformations up to 60% before rupturing, a value slightly higher than what we observed for *Hydra* tissue spheres. Our results are thus well in line with these observations, and both ask the question of the biological origin of the non-linear behavior of epithelial tissues which remains elusive for the moment.

Regarding the specific rheology of *Hydra*, our work is the first to focus specifically on regenerating tissue spheres but measurements exist on full adults. In (39), the authors employed a modified AFM experimental procedure to generate indentations on fixed, full *Hydras* with a 20μm wide sphere. By co-recording the deformation of the tissue and the force exerted, they observed a linear relationship between both which allowed them to estimate the elastic modulus of these samples. One key result was the observation that the elasticity of the body column was inhomogeneous, the tissue being roughly three times stiffer near the head than close to the foot. Although it is a well-known fact that different experimental approaches can lead to different estimations of rheological properties (53) and potential differences between fully mature *Hydras* and tissue spheres, our results are in good agreement with this work. Indeed, in the softest part of the adult, which we would imagine to most resemble tissue spheres, the authors report an elastic modulus on the order of 400Pa. Since the deformations involved are of a few microns, we expect these measurements to probe the small deformation regime where our own measurements also find an elastic modulus on the order of 400Pa. In the large deformation regime, our results also confirm those of (5) which, in order to properly reproduce the osmotic oscillations based on chemical considerations implemented an elastic modulus *E* = 11*kPa*. Still, AFM measurements on tissue spheres, albeit technically challenging, would be a great complement to our work to obtain a comprehensive understanding of the mechanics of these samples.

Another consequence of our focus on osmotic oscillations and tissue-scale mechanics is that our mechanical description of the tissue spheres assumes a homogenous shell, ignoring the complex three-layer (endoderm-extra-cellular matrix-ectoderm) structure of the samples. It is clearly possible that the deformations and stresses induced by the osmotic oscillations are differently transmitted and have different effects on these different layers, a refinement which will need to be addressed in the future. In particular, it was recently suggested that the mechanical properties of the ECM, which were not specifically measured here, could be linked to morphogenesis in *Hydra* (54).

Since we believe deformations and stresses within the tissues to be central in the morphogenesis of *Hydra*, another natural question is the existence of inhomogeneities of deformations, forces or rheological properties within the same sample. Mechanical fluctuations, if coupled to biochemistry, could well be sufficient to destabilize a reaction-diffusion system into a patterned state. Our own experimental setup could allow to measure fluctuations in rheological properties by repeating measurements on the same samples with small holes but at different locations, the main limitation being the lack of control of the position of the tested zone. In the same logic, it has been proposed that the amount of HyWnt3 protein could locally affect the Young’s modulus of the tissue (25) which could couple the chemical patterning of the head organizer and morphogenesis. If the future location of the head becomes softer, the homogenous osmotic pressure will deform this region more, leading to a deviation from a spherical shape to an oblong one. Here too, our setup could allow to measure the Young’s modulus of tissue spheres overexpressing HyWnt3 to directly test this hypothesis.

## Conclusion

In this work, we have employed an original parallelized microaspiration device to characterize the mechanical behavior of *Hydra* tissue spheres. Although we found different rheological behaviors depending on the applied stresses, the main result we obtained is that during normal osmotic oscillations, these samples behave largely as hyperelastic thin shells, displaying a nonlinear, stress-stiffening rheology. Doing so, we also obtained quantitative measurements of their relevant rheological parameters (*E* = (4.4 ± 3.3) . 10^2^ Pa, *η* = (2.4 ± 1.2) . 10^6^ Pa. s, critical applied pressures for viscous behavior and tissue rupture). This allowed us to estimate the stresses acting within the tissues and, in particular, the critical stress at which they start rupturing. These results, by allowing a quantification of both stresses and deformations during *Hydra* regeneration, will hopefully pave the way for quantitative approaches aiming at correlating these mechanical cues with molecular ones to understand *Hydra* patterning as an integrated mechano-biochemical process.

## Author contributions

OCE designed the research. ZBM, MD and PJ manufactured the inserts. TP, ABB, PM and OCE performed the research and analyzed data. TP, PM and OCE wrote the manuscript which was reviewed by all authors.

## Acknowledgments

This work was funded by the Mission pour les Initiatives Transverses et Interdisciplinaires of CNRS (Project MeChemReg to OCE and PM). We thank H. Delanoë-Ayari for fruitful discussions.

## References

1. Galliot, B., and V. Schmid. 2002. Cnidarians as a model system for understanding evolution and regeneration. Int. J. Dev. Biol. 46:39–48.

2. Technau, U., and R.E. Steele. 2011. Evolutionary crossroads in developmental biology: Cnidaria. Development. 138:1447–58.

3. Gierer, A., S. Berking, H. Bode, C.N. David, K. Flick, G. Hansmann, H. Schaller, and E. Trenkner. 1972. Regeneration of hydra from reaggregated cells. Nat. New Biol. 239:98–101.

4. Cochet-Escartin, O., T.T. Locke, W.H. Shi, R.E. Steele, and E.-M.S. Collins. 2017. Physical Mechanisms Driving Cell Sorting in Hydra. Biophys. J. 113:2827–2841.

5. Kücken, M., J. Soriano, P.A. Pullarkat, A. Ott, and E.M. Nicola. 2008. An osmoregulatory basis for shape oscillations in regenerating hydra. Biophys. J. 95:978–85.

6. Fütterer, C., C. Colombo, F. Jülicher, and A. Ott. 2003. Morphogenetic oscillations during symmetry breaking of regenerating Hydra vulgaris cells. Europhys. Lett. 64:137.

7. Broun, M., L. Gee, B. Reinhardt, and H.R. Bode. 2005. Formation of the head organizer in hydra involves the canonical Wnt pathway. Development. 132.

8. Bode, H.R. 2009. Axial Patterning in Hydra. Cold Spring Harb. Perspect. Biol. 1.

9. Böttger, A., and M. Hassel. 2012. Hydra, a model system to trace the emergence of boundaries in developing eumetazoans. Int. J. Dev. Biol. 56:583–91.

10. Soriano, J., C. Colombo, and A. Ott. 2006. Hydra molecular network reaches criticality at the symmetry-breaking axis-defining moment. Phys. Rev. Lett. 97:258102.

11. Turing, A.M. 1952. The Chemical Basis of Morphogenesis. Philos. Trans. R. Soc. B Biol. Sci. 237:37–72.

12. Marcon, L., and J. Sharpe. 2012. Turing patterns in development: What about the horse part? Curr. Opin. Genet. Dev. 22:578–584.

13. Schweisguth, F., and F. Corson. 2019. Self-Organization in Pattern Formation. Dev. Cell. 49:659–677.

14. Gierer, A., and H. Meinhardt. 1972. A theory of biological pattern formation. Kybernetik. 12:30–9.

15. Nakamura, Y., C.D. Tsiairis, S. Özbek, and T.W. Holstein. 2011. Autoregulatory and repressive inputs localize Hydra Wnt3 to the head organizer. Proc. Natl. Acad. Sci. U. S. A. 108:9137–42.

16. Augustin, R., A. Franke, K. Khalturin, R. Kiko, S. Siebert, G. Hemmrich, and T.C.G. Bosch. 2006. Dickkopf related genes are components of the positional value gradient in Hydra. Dev. Biol. 296:62–70.

17. Vogg, M.C., L. Beccari, L. Iglesias Ollé, C. Rampon, S. Vriz, C. Perruchoud, Y. Wenger, and B. Galliot. 2019. An evolutionarily-conserved Wnt3/β-catenin/Sp5 feedback loop restricts head organizer activity in Hydra. Nat. Commun. 10:312.

18. Ziegler, B., I. Yiallouros, B. Trageser, S. Kumar, M. Mercker, S. Kling, M. Fath, U. Warnken, M. Schnölzer, T.W. Holstein, M. Hartl, A. Marciniak-Czochra, J. Stetefeld, W. Stöcker, and S. Özbek. 2021. The Wnt-specific astacin proteinase HAS-7 restricts head organizer formation in Hydra. BMC Biol. 19:1–22.

19. Soriano, J., S. Rüdiger, P. Pullarkat, and A. Ott. 2009. Mechanogenetic coupling of Hydra symmetry breaking and driven Turing instability model. Biophys. J. 96:1649–60.

20. Ferenc, J., P. Papasaikas, J. Ferralli, Y. Nakamura, S. Smallwood, and C.D. Tsiairis. 2021. Mechanical oscillations orchestrate axial patterning through Wnt activation in Hydra. Sci. Adv. 7:6897.

21. Desprat, N., W. Supatto, P.-A. Pouille, E. Beaurepaire, and E. Farge. 2008. Tissue deformation modulates twist expression to determine anterior midgut differentiation in Drosophila embryos. Dev. Cell. 15:470–7.

22. Gudipaty, S.A., J. Lindblom, P.D. Loftus, M.J. Redd, K. Edes, C.F. Davey, V. Krishnegowda, and J. Rosenblatt. 2017. Mechanical stretch triggers rapid epithelial cell division through Piezo1. Nature. 543:118–121.

23. Pouille, P.-A., P. Ahmadi, A.-C. Brunet, and E. Farge. 2009. Mechanical signals trigger Myosin II redistribution and mesoderm invagination in Drosophila embryos. Sci. Signal. 2:ra16.

24. Shinozuka, T., R. Takada, S. Yoshida, S. Yonemura, and S. Takada. 2019. Wnt produced by stretched roof-plate cells is required for the promotion of cell proliferation around the central canal of the spinal cord. Dev. 146.

25. Mercker, M., A. Köthe, and A. Marciniak-Czochra. 2015. Mechanochemical symmetry breaking in Hydra aggregates. Biophys. J. 108:2396–407.

26. Brinkmann, F., M. Mercker, T. Richter, and A. Marciniak-Czochra. 2018. Post-Turing tissue pattern formation: Advent of mechanochemistry. PLoS Comput. Biol. 14:e1006259.

27. Wang, R., T. Goel, K. Khazoyan, Z. Sabry, H.J. Quan, P.H. Diamond, and E.-M.S. Collins. 2019. Mouth Function Determines the Shape Oscillation Pattern in Regenerating Hydra Tissue Spheres. Biophys. J. 117:1145–1155.

28. Livshits, A., L. Shani-Zerbib, Y. Maroudas-Sacks, E. Braun, and K. Keren. 2017. Structural Inheritance of the Actin Cytoskeletal Organization Determines the Body Axis in Regenerating Hydra. Cell Rep. 18:1410–1421.

29. Wang, R., R.E. Steele, and E.M.S. Collins. 2020. Wnt signaling determines body axis polarity in regenerating Hydra tissue fragments. Dev. Biol. 467:88–94.

30. Wang, R., A.L. Bialas, T. Goel, and E.M.S. Collins. 2023. Mechano-Chemical Coupling in Hydra Regeneration and Patterning. Integr. Comp. Biol. 63:1422–1441.

31. Veschgini, M., F. Gebert, N. Khangai, H. Ito, R. Suzuki, T.W. Holstein, Y. Mae, T. Arai, and M. Tanaka. 2016. Tracking mechanical and morphological dynamics of regenerating Hydra tissue fragments using a two fingered micro-robotic hand. Appl. Phys. Lett. 108.

32. Evans, E.A. 1973. New Membrane Concept Applied to the Analysis of Fluid Shear- and Micropipette-Deformed Red Blood Cells. Biophys. J. 13:941–954.

33. Guevorkian, K., M.-J. Colbert, M. Durth, S. Dufour, and F. Brochard-Wyart. 2010. Aspiration of Biological Viscoelastic Drops. Phys. Rev. Lett. 104:1–4.

34. Hochmuth, R.M. 2000. Micropipette aspiration of living cells. J. Biomech. 33:15–22.

35. Lim, C.T., E.H. Zhou, and S.T. Quek. 2006. Mechanical models for living cells--a review. J. Biomech. 39:195–216.

36. Landiech, S., M. Elias, P. Lapèze, H. Ajiyel, M. Plancke, A. Laborde, F. Mesnilgrente, D. Bourrier, D. Berti, C. Montis, L. Mazenq, J. Baldo, C. Roux, M. Delarue, and P. Joseph. 2023. Parallel on-chip micropipettes enabling quantitative multiplexed characterization of vesicle mechanics and cell aggregates rheology. bioRxiv. 2023.10.19.562871.

37. Holzapfel, G.A. 2000. Nonlinear solid mechanics: a continuum approach for engineering science. Wiley, Hoboken.

38. Green, A.E., and W. Zerna. 1968. Theoretical Elasticity. Dover Publications, Mineola.

39. Naik, S., M. Unni, D. Sinha, S.S. Rajput, P.C. Reddy, E. Kartvelishvily, I. Solomonov, I. Sagi, A. Chatterji, S. Patil, and S. Galande. 2020. Differential tissue stiffness of body column facilitates locomotion of Hydra on solid substrates. J. Exp. Biol. 223.

40. Boot, R.C., A. Roscani, L. van Buren, S. Maity, G.H. Koenderink, and P.E. Boukany. 2023. High-throughput mechanophenotyping of multicellular spheroids using a microfluidic micropipette aspiration chip. Lab Chip. 23:1768.

41. Guevorkian, K., F. Brochard-Wyart, and D. Gonzalez-Rodriguez. 2021. Flow dynamics of 3D multicellular systems into capillaries. In: Viscoelasticity and Collective Cell Migration: An Interdisciplinary Perspective Across Levels of Organization. Academic Press. pp. 193–223.

42. Harris, A.R., L. Peter, J. Bellis, B. Baum, A.J. Kabla, and G.T. Charras. 2012. Characterizing the mechanics of cultured cell monolayers. Proc. Natl. Acad. Sci. U. S. A. 109:16449–16454.

43. Brinkmann, F. 2020. Mathematical models and numerical simulation of mechanochemical pattern formation in biological tissues. PhD thesis, Ruprecht-Karls-Universität.

44. Ugural, A.C. 2009. Stresses in beams, plates, and shells, third edition. CRC Press, Boca Raton.

45. Takeichi, M. 1977. Functional correlation between cell adhesive properties and some cell surface proteins. J. Cell Biol. 75:464–474.

46. Straight, A.F., A. Cheung, J. Limouze, I. Chen, N.J. Westwood, J.R. Sellers, and T.J. Mitchison. 2003. Dissecting temporal and spatial control of cytokinesis with a myosin II inhibitor. Science (80-.). 299:1743–1747.

47. Sato, M., D.P. Theret, L.T. Wheeler, N. Ohshima, and R.M. Nerem. 1990. Application of the Micropipette Technique to the Measurement of Cultured Porcine Aortic Endothelial Cell Viscoelastic Properties. J. Biomech. Eng. 112:263–268.

48. Tsai, M.A., R.S. Frank, and R.E. Waugh. 1993. Passive mechanical behavior of human neutrophils: power-law fluid. Biophys. J. 65:2078–2088.

49. Ruiz-Herrero, T., K. Alessandri, B. V. Gurchenkov, P. Nassoy, and L. Mahadevan. 2017. Organ size control via hydraulically gated oscillations. Development. 144:4422–4427.

50. Lien, J.C., and Y. li Wang. 2021. Cyclic stretching-induced epithelial cell reorientation is driven by microtubule-modulated transverse extension during the relaxation phase. Sci. Reports 2021 111. 11:1–12.

51. Gérémie, L., E. Ilker, M. Bernheim-Dennery, C. Cavaniol, J.L. Viovy, D.M. Vignjevic, J.F. Joanny, and S. Descroix. 2022. Evolution of a confluent gut epithelium under on-chip cyclic stretching. Phys. Rev. Res. 4:023032.

52. Fink, J., N. Carpi, T. Betz, A. Bétard, M. Chebah, A. Azioune, M. Bornens, C. Sykes, L. Fetler, D. Cuvelier, and M. Piel. 2011. External forces control mitotic spindle positioning. Nat. Cell Biol. 2011 137. 13:771–778.

53. Wu, P.H., D.R. Ben Aroush, A. Asnacios, W.C. Chen, M.E. Dokukin, B.L. Doss, P. Durand-Smet, A. Ekpenyong, J. Guck, N. V. Guz, P.A. Janmey, J.S.H. Lee, N.M. Moore, A. Ott, Y.C. Poh, R. Ros, M. Sander, I. Sokolov, J.R. Staunton, N. Wang, G. Whyte, and D. Wirtz. 2018. A comparison of methods to assess cell mechanical properties. Nat. Methods 2018 157. 15:491–498.

54. Veschgini, M., R. Suzuki, S. Kling, H.O. Petersen, B.G. Bergheim, W. Abuillan, P. Linke, S. Kaufmann, M. Burghammer, U. Engel, F. Stein, S. Özbek, T.W. Holstein, and M. Tanaka. 2023. Wnt/β-catenin signaling induces axial elasticity patterns of Hydra extracellular matrix. iScience. 26:106416.

