## Supplementary Information for "Mechanical characterization of regenerating *Hydra* tissue spheres"

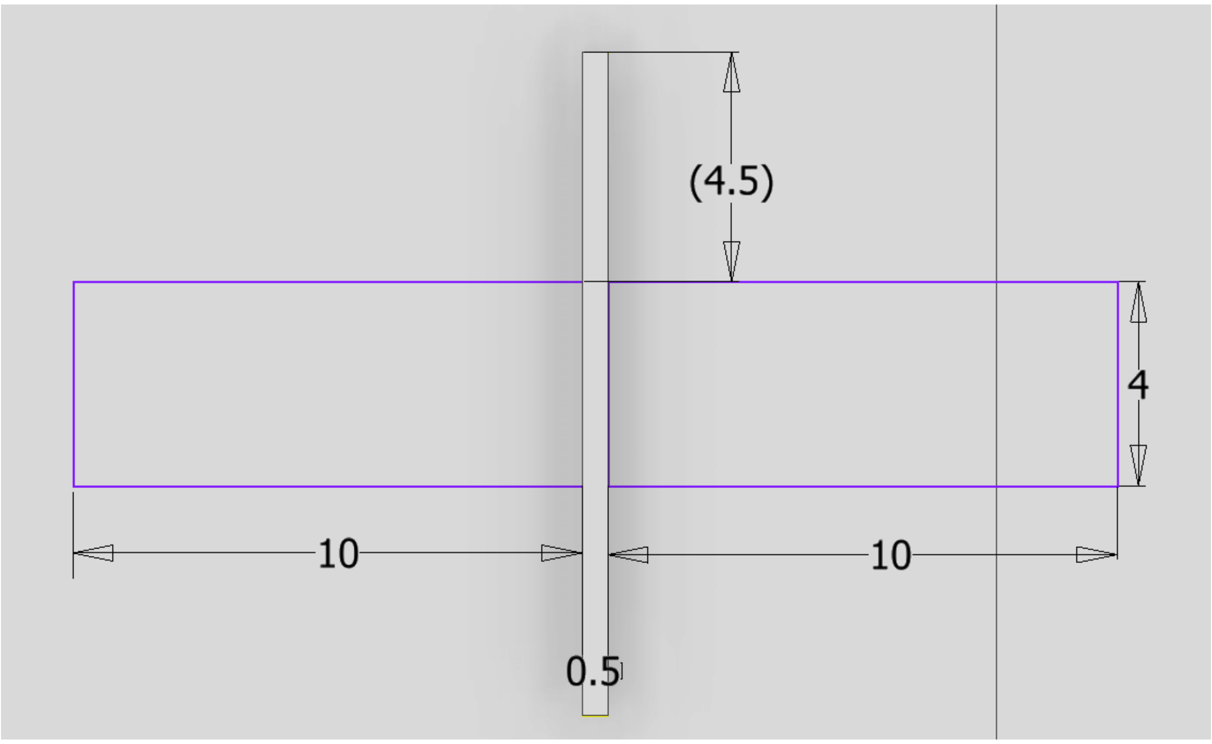


Fig S1. Schematics and dimensions (in mm) of the chamber used in micro aspiration experiments.

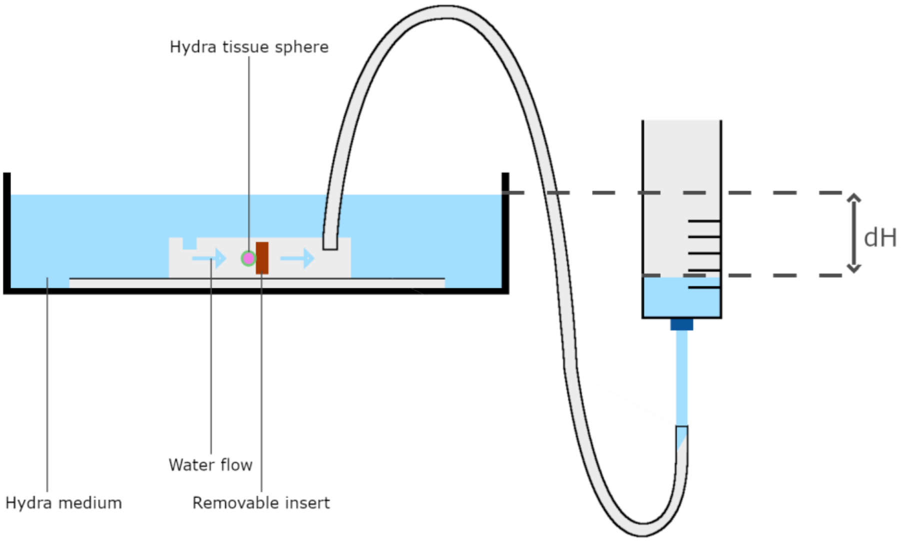


Fig S2. Schematics of the micro-aspiration full setup.

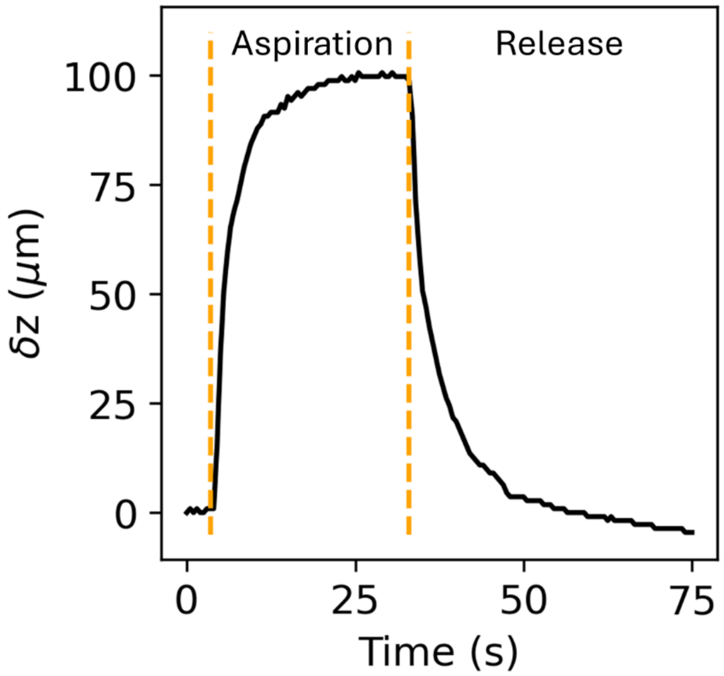


Fig S3. Reversible dynamics of aspirated length during aspiration and release (shown by orange dashed lines) at $\Delta P = 1500Pa$ in a square opening.

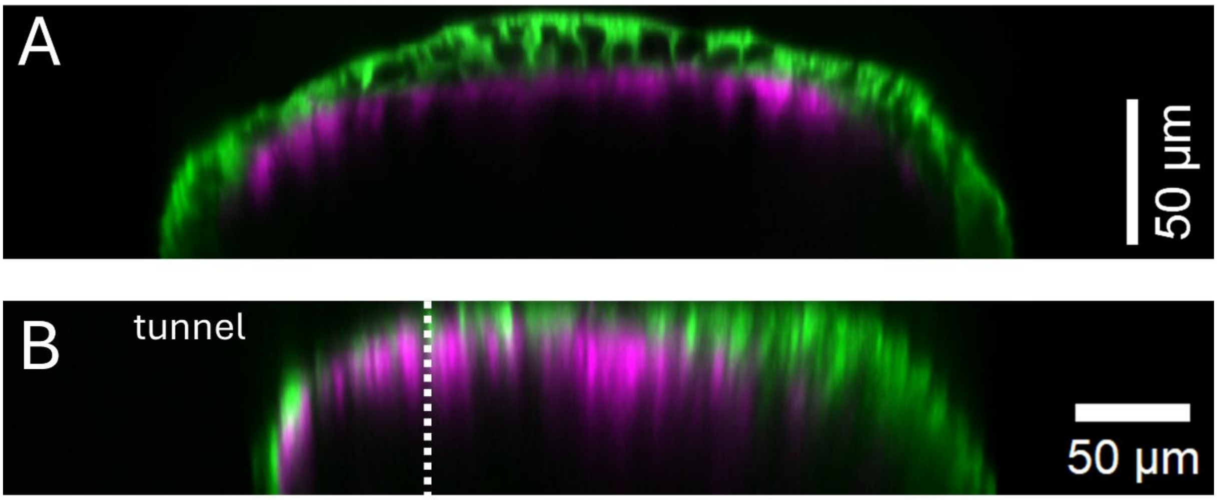


Fig S4. A: Side view of a “watermelon” tissue piece (ectoderm in green, endoderm in magenta) acquired by spinning disk microscopy. B: Similar side view of a watermelon tissue piece aspirated in a square tunnel. The entrance of the tunnel is visualized as a dashed white line. The difference in resolution between the two images is due to the optical properties of the microfluidic device, this is more visible within the tunnel. No measurable changes in thickness were observed.

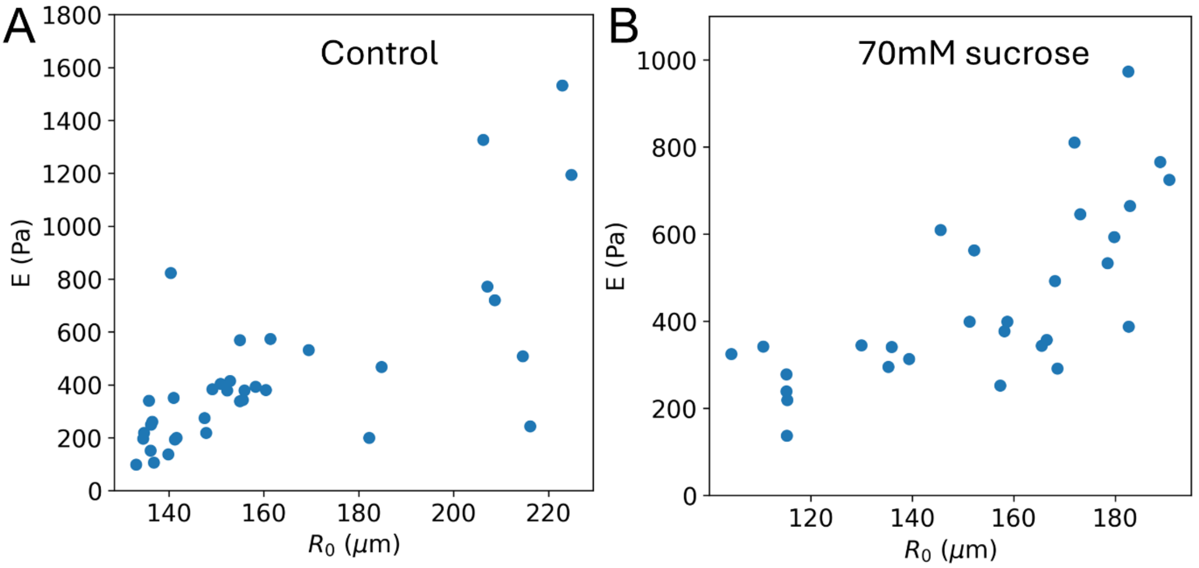


Fig S5. Plot of Young's modulus as a function of initial tissue sphere ratio under normal conditions and 70mM sucrose.

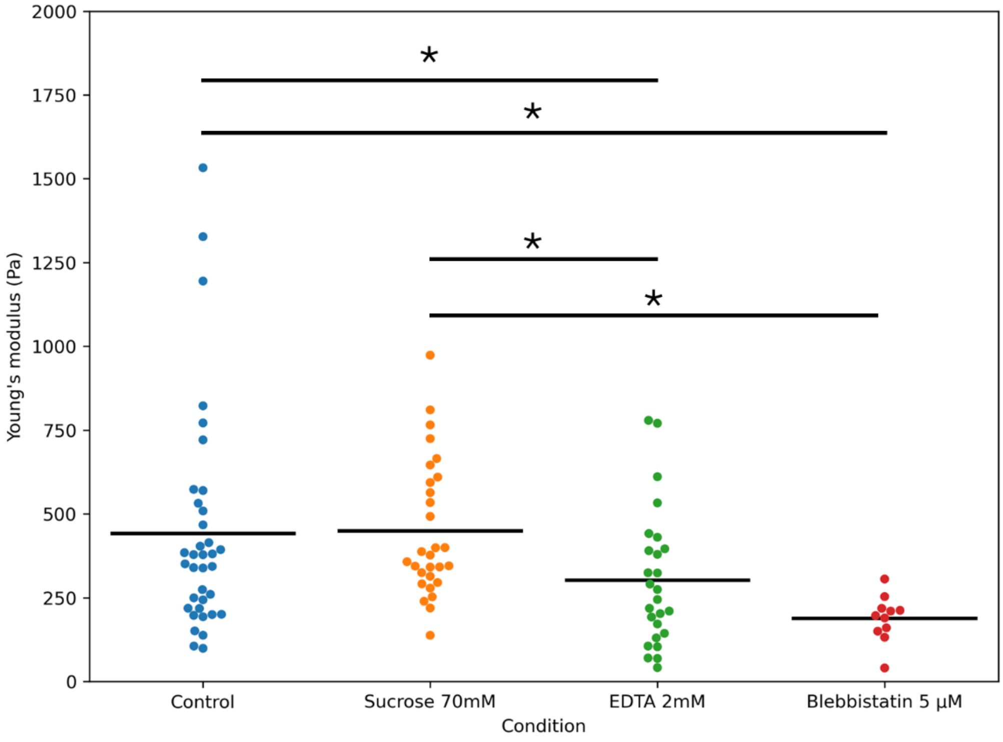


Fig S6. Measurement of Young's modulus under different chemical modulations. The black lines represent the mean values. Statistically significant differences were estimated using an independent two-sample t-test ($p<0.05$) and are represented by stars. Control: $E = (4.4 \pm 3.3) \,.\, 10^2\ Pa$, sucrose: $E = (4.5 \pm 2.0) \,.\, 10^2\ Pa$, EDTA: $E = (3.0 \pm 2.0) \,.\, 10^2\ Pa$, blebbistatin: $E = (1.9 \pm 0.7) \,.\, 10^2\ Pa$. mean ± standard deviation, n=36, 29, 26 and 11, respectively.

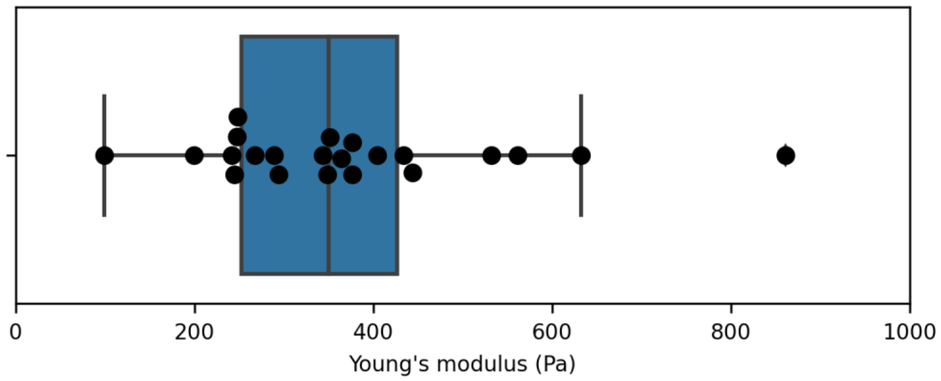


Fig S7. Estimation of Young's modulus in the visco-elastic phase, n=22. We find, here, $E = (3.7 \pm 1.6)\,.\,10^2\ Pa$, consistent with measurements in the elastic phase.

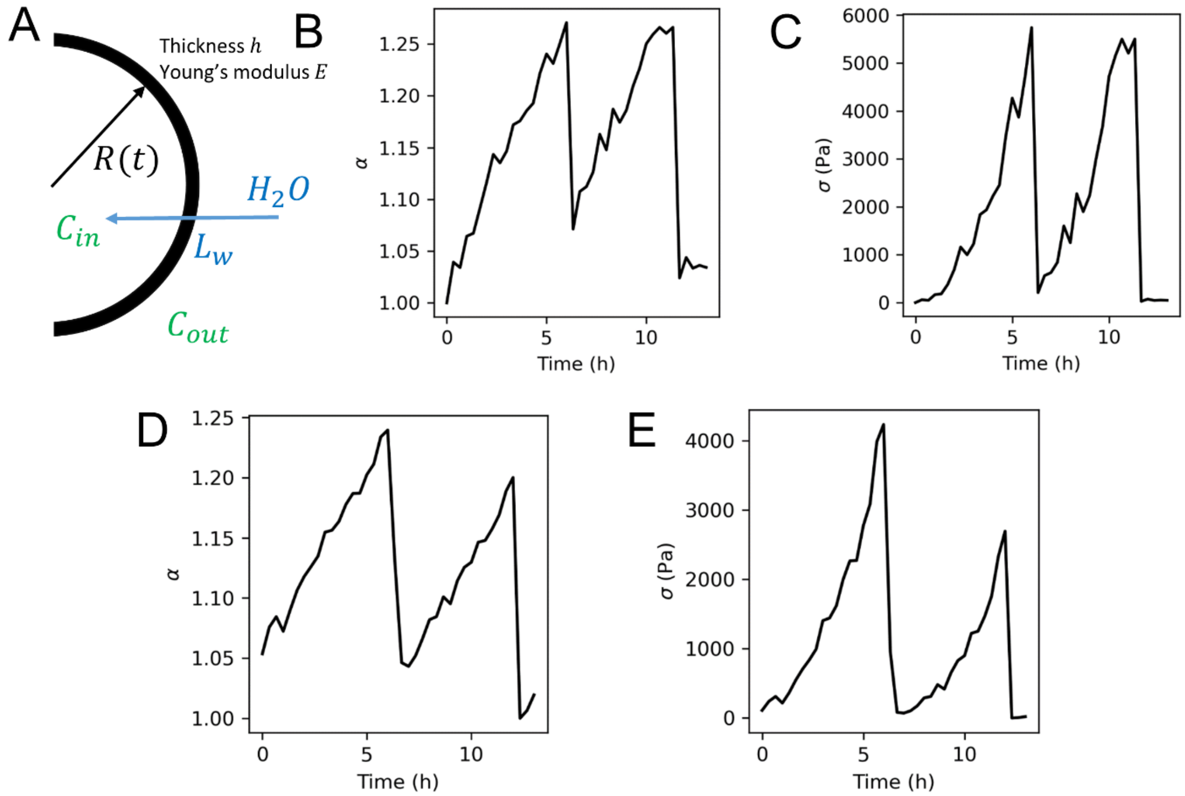


Fig S8. A: schematics and notations used to describe the mechanics of the tissue sphere. B-D: Two examples of osmotic oscillations displayed as the stretch ratio $\alpha = \frac{R(t)}{R_0}$ as a function of time. C-E: resulting dynamics of the hoop stress $\sigma$ as a function of time using Eq 11 and $\mu = 148Pa,\ \lambda = 100\mu$ (see Main text).
